# The impact of contingency awareness on the neurocircuitry underlying pain-related fear and safety learning

**DOI:** 10.1101/2025.06.25.661520

**Authors:** Franziska Labrenz, Sigrid Elsenbruch, Adriane Icenhour

**Affiliations:** Department of Affective Neuroscience, Ruhr University Bochum, Bochum, Germany; Department of Medical Psychology and Medical Sociology, Ruhr University Bochum, Bochum, Germany; Department of Neurology, Center for Translational Neuro- and Behavioral Sciences, University Hospital Essen, University of Duisburg-Essen, Essen, Germany

**Keywords:** Contingency awareness, fMRI, interoceptive conditioning, pain-related fear conditioning, safety learning, visceral pain

## Abstract

Visceral pain-related fear, shaped by associative learning, drives maladaptive emotional reactions and may contribute to the chronicity of pain in disorders of gut-brain interaction. However, the role of contingency awareness remains unclear. In a translational model of pain-related conditioning, we investigated the brain-behavior relationships underlying contingency awareness in shaping the neural circuitry involved in visceral pain-related fear and safety learning.

Data from 75 healthy individuals undergoing differential conditioning were acquired in two functional magnetic resonance imaging studies. Visceral pain as unconditioned stimulus (US) was paired with a visual cue as conditioned stimulus (CS^+^) while another cue (CS^-^) remained unpaired. Differential neural responses to predictive cues were analyzed using a full factorial model and regression analyses to evaluate the predictive value of neural activation patterns based on contingency awareness.

Analyses revealed a significant interaction between CS-type and contingency awareness involving dorsolateral prefrontal cortex (dlPFC) and parahippocampus, driven by an enhanced CS^+^>CS^-^ differentiation in highly aware participants. The reverse contrast revealed widespread activation in fronto-parietal and limbic networks, more pronounced in the highly aware group. Regression analyses showed that enhanced CS^-^-related were associated with increased contingency awareness and CS^-^ valence change, while no activation clusters predictive of behavioral responses were found for CS^+^.

The recruitment of emotional arousal and executive control networks as a function of contingency awareness highlights its relevance in shaping pain- and, particularly, safety-predictive cue properties. These results suggest distinct processes for fear acquisition and inhibition, with significant implications for exposure-based treatments of disorders of gut-brain interaction.

## 1. Introduction

Associative learning and memory processes fundamentally shape intricate relationships between the brain and behavior, affecting a wide range of emotional, attentional, and executive processes (Bouton, 2018; Franklin & Grossberg, 2017). One of the most potent drivers of associative learning is fear, evoking a cascade of responses that manifest at behavioral, physiological, and neuronal levels (LeDoux, 2014; Maren, 2001).

Governed by the principles of Pavlovian conditioning, associative learning occurs when an initially neutral stimulus (conditioned stimulus, CS^+^) is repeatedly paired with a biologically relevant event (unconditioned stimulus, US), evoking a conditioned response (CR) upon subsequent presentation of the CS^+^ alone. This learning process is further influenced by conditioned inhibition through safety signal learning, which involves a CS^-^ predicting the absence of imminent threat (Christianson et al., 2012; Laing et al., 2021). The processes underlying fear conditioning are often assumed to be simple and automatic such that threatening stimuli forge stronger aversive associations, activate fear-related neural circuits without prior cognitive evaluation, and operate largely outside the scope of conscious control (Mineka & Öhman, 2002). Several studies support this view, suggesting that fear learning can occur without conscious awareness of the CS^+^-US relationship, meaning individuals do not necessarily need to recognize that the CS^+^ predicts the US in order to develop fear responses (Knight et al., 2009, 2006; Öhman & Mineka, 2001; Raio et al., 2012; Schultz & Helmstetter, 2010; Sevenster et al., 2014; Weike et al., 2007). On the other hand, there is also evidence suggesting that contingency awareness about the association between the CS^+^ and US is a prerequisite for successful fear conditioning (Klucken et al., 2009; Lovibond & Shanks, 2002; Mertens & Engelhard, 2020; Mitchell et al., 2009; Tabbert et al., 2011, 2006). These conflicting findings have sparked a burgeoning interest in understanding the putative role of contingency awareness in fear learning, and recent advances in behavioral neurosciences have motivated studies to explore the brain-behavior relationships that underlie this process.

Several studies have reported enhanced threat-induced neural activation involving prefrontal areas, the anterior cingulate cortex, insula, amygdala, hippocampus, and parietal regions (Baeuchl et al., 2019; Carter et al., 2006; Klucken et al., 2009; Madaboosi et al., 2021; Tabbert et al., 2011) in contingency aware, but not unaware, participants. The involvement of these areas as essential components of the salience and central executive networks suggests that the integration, cognitive modulation, and memory formation of fear-related cue properties may be of crucial relevance for acquiring contingency awareness. Conversely, increased neural responses in unaware compared to aware participants involving partly overlapping regions including prefrontal and cingulate cortices, parietal lobule as well as the insula, amygdala, and hippocampus appear to contradict these observations (Klucken et al., 2009; Lam et al., 2023; Tabbert et al., 2011, 2006).

The heterogeneity of these results may stem from the influence of moderating variables, such as outcome measures, procedural aspects, or stimulus material (Mertens & Engelhard, 2020). Accumulating evidence suggests a particular impact of the biological salience and ecological validity of the applied US on learning efficacy, with pain, as a hard-wired signal of bodily harm, demonstrably constituting one of the strongest motivators for learning (Meulders, 2020). Previous studies have documented that interoceptive visceral pain is perceived as more unpleasant and fear-provoking compared to other pain modalities (Dunckley et al., 2005; Koenen et al., 2018, 2017), and that associations related to visceral pain are preferentially learned, stored, and remembered (Koenen et al., 2021). Our work on visceral pain-related conditioning has previously expanded on these findings, showing that experimental visceral pain induces robust learned fear responses to pain-predictive cues, involving enhanced recruitment of key regions of the central fear, salience, and executive control networks (Gramsch et al., 2014; Icenhour et al., 2021, 2017, 2015a; Kattoor et al., 2013; Labrenz et al., 2016). In the presence of such a highly aversive threat, cues signaling the absence of imminent pain acquire properties of relief and become potent inhibitors of fear (Christianson et al., 2012; Laing et al., 2021; Odriozola & Gee, 2021) highlighting the distinct relevance of safety learning and memory processes in the context of visceral pain (Labrenz et al., 2022a, 2015a). However, the specific role of accurate contingency awareness in shaping threat- and safety-related responses in the context of visceral pain and the neural underpinnings of these processes remain largely unexplored.

Providing first evidence, an analysis of behavioral data from a large, pooled sample of healthy volunteers as included herein, revealed that contingency awareness shapes the acquisition and extinction of emotional responses associated with visceral pain (Labrenz et al., 2015a). Specifically, individuals showing accurate awareness of cue-pain contingencies demonstrated a stronger differentiation in emotional valence between learned threat and safety cues after acquisition and extinction. Moreover, contingency awareness was found to predict the valence of the safety, but not the threat cue, indicating that the acquisition of positive emotional responses may particularly require accurate awareness. These findings suggest distinct mechanisms underlying the acquisition of conditioned responses to threat and safety cues, at least at the behavioral level. However, further evaluation at the neural level is needed.

The aim of this study was therefore to elucidate the brain-behavior relationships underlying contingency awareness in shaping the neural circuitry involved in visceral pain-related fear and safety learning, based on neuroimaging data from two independent studies. Building upon our behavioral findings and existing fMRI evidence from the broader fear conditioning literature, we hypothesized that contingency awareness would shape distinct neural responses to threat- and safety-predictive cues. Specifically, we expected that enhanced involvement of fear-related limbic regions in response to the CS^+^, and increased engagement of prefrontal and parietal areas in response to the CS^-^, would reflect heightened working memory load, attentional processing, and regulatory control. Enhanced recruitment of key regions within the salience, central executive, and fear networks was anticipated in participants who accurately reported cue-pain contingencies.

## 2. Materials and Methods

### 2.1 Participants

For the current re-analysis, functional neuroimaging data from 75 healthy volunteers were pooled from two independent studies addressing visceral, pain-related conditioning (Icenhour et al., 2015a; Labrenz et al., 2016). Behavioral data from this sample were previously published (Labrenz et al., 2015a). Recruitment procedures, as well as inclusion and exclusion data, were identical for both studies. The inclusion criteria for study participation were as follows: age between 18 and 60 years, body mass index (BMI) between 18 and 30 kg/m², no history of any medical or psychiatric condition, and no chronic medication use, except for hormonal contraceptives or occasional over-the-counter allergy or pain medications, as reported by the participant. Symptoms suggestive of functional or gastrointestinal conditions were excluded based on a standardized in-house questionnaire (Lacourt et al., 2014). All participants underwent a physical examination to exclude perianal tissue damage (e.g., fissures or painful hemorrhoids) that could interfere with balloon placement. In women, pregnancy was ruled out using a commercially available urinary test on the study day. Current symptoms of anxiety and depression were assessed with the German version of the Hospital Anxiety and Depression Scale (HADS) using the published, clinically relevant cut-off values of ≥ 8 (Herrmann-Lingen et al., 2005). The study protocols adhered to the guidelines outlined in the Declaration of Helsinki and were approved by the local Ethics Committee of the University Hospital Essen, Germany (protocol number 10-4493). All participants provided written informed consent and were reimbursed for their participation.

### 2.2 Study designs and procedures

The study design and procedures were identical for both studies and have been previously described in detail (Icenhour et al., 2015a; Labrenz et al., 2016). Briefly, sensory and pain thresholds for rectal distensions, used as an experimental visceral pain model, were assessed for each participant to determine individual pressures for aversive visceral US applied in the experimental task. A structural MRI scan was completed, followed by fMRI measurements, during which a differential fear conditioning paradigm was conducted. In this paradigm, one visual cue (CS^+^) was repeatedly paired with a rectal distension (US), while another visual cue (CS^-^) was never followed by the US. Overall, 16 CS^+^, of which 12 CS^+^ were followed by the US and 16 CS^-^ were presented in a pseudo-randomized order. The duration of both CSs varied between 8 and 12 s until the onset of the US, which lasted 16.8 s. Inter-stimulus intervals were 20 s. Following conditioning, US intensity ratings were assessed using online visual analog scales (VAS) with an MRI-compatible hand-held fiber optic response system (LUMItouch™, Photon Control Inc., Burnaby, BC, Canada). Participants were prompted to respond to the question, “How painful did you perceive the distensions?” with VAS endpoints labeled “not painful at all” (0) and “very painful” (100). Additionally, CS valence as a behavioral indicator of emotional learning was assessed by asking participants, “How do you perceive the circle/ square?”. Responses were logged on VAS with end points “very unpleasant” (−100) to “very pleasant” (+100). Behavioral data from this pooled sample (Labrenz et al., 2015a) and from the respective subsamples (Icenhour et al., 2015a; Labrenz et al., 2016) have previously been published.

Following visceral pain-related conditioning, CS-US contingency ratings were assessed to estimate participants’ awareness of CS-US pairings, which was, in fact, 75 %. Participants were prompted to respond to the question “How often was the circle/ square followed by a rectal distension?” using a VAS with endpoints labeled “never” (0) and “always” (100). These ratings were further transformed into an integrative contingency accuracy score (in %) to quantify awareness of the actual CS-US reinforcement rates adequately and for group allocation, as previously established and described in detail (Labrenz et al., 2015a). This measure was chosen because it quantifies the distinct influences of pain- and safety-related contingencies beyond mere differentiation and provides an explicit measure of accuracy in contingency estimations within the partial reinforcement schedule applied. Subgroups with high versus low contingency accuracy were defined based on a median split, resulting in two groups: one with a high mean accuracy (N = 41) and the other with low accuracy (N = 34).

### 2.3 Analyses of sociodemographic and behavioral data

Sociodemographics, psychological measures, and thresholds, as well as changes in CS valence, operationalized as 𝚫 valence post-acquisition – baseline and contingency accuracy scores in % as behavioral data implemented in the current fMRI analyses, were analyzed using IBM SPSS Statistics 27.0 (IBM Corporation, Armonk, NY, USA). Initially, normal distribution was confirmed using Kolmogorov–Smirnov tests, and two-sided parametric testing was subsequently applied. Descriptive statistics are provided as mean ± standard error of the mean (SEM), and results of independent samples t-tests comparing the low and high contingency awareness group are given with significance defined as *p* < .05.

### 2.4 MRI data acquisition and analyses

All MR images were acquired on a whole-body 3T scanner (MAGNETOM Skyra, Siemens Healthcare, Erlangen, Germany) using a 32-channel head coil. Structural images were acquired using a T1-weighted 3D-MPRAGE sequence with the following parameters: TR 1900 ms, TE 2.13 ms, flip angle 9°, FOV 239 × 239 mm², 192 slices, slice thickness 0.9 mm, voxel size 0.9 × 0.9 × 0.9 mm³, matrix 256 × 256, and GRAPPA *r* = 2. Functional images were acquired using a multi-echo EPI sequence with three echoes (echo 1 TE 13.0 ms, echo 2 TE 28.9 ms, echo 3 TE 44.8 ms) (Poser et al., 2006) and TR 2000 ms, flip angle 90°, FOV 220 × 220 mm2, matrix 80 × 80 mm2, GRAPPA *r* = 3 with 36 transversal slices angulated in the direction of the corpus callosum, slice thickness of 3 mm, slice gap 0.6 mm and a voxel size of 2.8 × 2.8 × 3.0 mm.

Functional MR images were analyzed using SPM12 (Wellcome Trust Centre for Neuroimaging, UCL, London, UK; RRID: SCR_007037) implemented in MATLAB R2019b (MathWorks Inc., Sherborn, MA, USA; RRID: SCR_001622). Initially, the three echoes of the multi-echo EPI sequence were combined automatically (Poser et al., 2006) and preprocessed, involving motion correction, normalization to a standardized template provided by the Montreal Neurological Institute, and smoothing with an isotropic Gaussian kernel of 8 mm. To correct for low frequency drifts, a temporal high-pass filter with a cutoff at 128 s was applied and serial autocorrelations were considered utilizing an autoregressive model first-order correction.

First-level analyses included applying a general linear model to the preprocessed EPI images. For each voxel, the time series was fitted with a corresponding task regressor modeling a boxcar convolved with the canonical hemodynamic response function. The regressors included CS^+^ with 16 trials, CS^-^ with 16 trials, and US with 12 trials. Each regressor was modeled with the respective stimulus onset and the entire duration of stimulus presentation of 8-12 s for the CS and 16.8 s for the US. Additionally, six realignment parameters for translation and rotation were entered as nuisance regressors to account for the rigid body transformation.

The ensuing first-level contrasts for CS^+^ and CS^-^ were computed and entered into second-level group analyses using a full factorial model with group (high contingency awareness, low contingency awareness) and CS (CS^+^, CS^-^) as factors. Within this model, we first assessed the interaction effect between group and CS. Post-hoc paired t-tests were computed to assess the differential neural activation in response to CS for the high and low contingency awareness groups, resulting in the contrasts: High awareness CS^+^ > CS^-^, Low awareness CS^+^ > CS^-^, High awareness CS^-^ > CS^+^ and Low awareness CS^-^ > CS^+^.

Regression analyses were further performed to assess the modulation of neural activation patterns as a function of contingency awareness and changes in CS valence. To test for effects of interest in the full sample, changes in CS valence from baseline to post-conditioning for each CS separately and contingency accuracy scores were entered as covariates. For significant results, post hoc tests for each covariate of interest were computed to discern the directionality of effects. Finally, exploratory analyses were conducted to exclude putative group differences in pain processing induced by painful visceral US.

All second-level analyses were assessed in *a priori* defined regions-of-interest (ROI) based on results of previous fMRI studies on the role of contingency awareness in classical fear conditioning (Baeuchl et al., 2019; Carter et al., 2006; Klucken et al., 2009; Madaboosi et al., 2021; Tabbert et al., 2011) and on visceral pain-related fear learning and extinction (Benson et al., 2014; Elsenbruch et al., 2014; Icenhour et al., 2021, 2015a, b; Koenen et al., 2021, 2018). ROI included the dorsolateral (dlPFC), ventrolateral (vlPFC), ventromedial prefrontal (vmPFC), orbitofrontal (OFC), and cingulate cortices (ACC, MCC, PCC), insula, inferior (IPL), medial (precuneus), and superior parietal lobules (SPL), amygdala, hippocampus and parahippocampus. All ROI analyses were performed using anatomical templates constructed from the WFU Pick Atlas toolbox (Version 3.5) (Maldjian et al., 2003), as implemented in SPM. A cluster-level extent threshold controlling for family-wise error rates (FWE) within the respective research area was applied, and results are reported with peak MNI coordinates and exact unilateral *p* values (all *p*_FWE_ < .05).

## 3. Results

### 3.1 Sample characterization

A characterization of the full sample and a comparison between groups with high and low contingency awareness with respect to sociodemographics, psychological variables, sensory and pain thresholds, as well as behavioral correlates of associative learning are summarized in Table 1. Overall, the 75 young and healthy volunteers had a BMI within the normal weight range and low scores on anxiety and depression, well below the clinical cut-off score. Visceral sensory and pain thresholds determined during the calibration procedure were comparable to those in previous samples of young and healthy volunteers (Icenhour et al., 2021, 2019; Koenen et al., 2021, 2017; Labrenz et al., 2016).

**Table 1.** Characterization of the full sample and contingency awareness groups.

|  | Full sample | High CA | Low CA | t-statistic |
| --- | --- | --- | --- | --- |
| Age (years) | 28.87 ± 1.11 | 27.07 ± 1.14 | 31.03 ± 1.98 | 1.73 |
| BMI (kg/m<sup>2</sup>) | 23.78 ± 0.49 | 22.32 ± 0.48 | 25.54 ± 0.82 | 3.40** |
| HADS Anxiety | 2.84 ± 0.28 | 3.15 ± 0.43 | 2.47 ± 0.32 | 1.26 |
| HADS Depression | 1.25 ± 0.18 | 1.61 ± 0.29 | 0.82 ± 0.17 | 2.30* |
| Sensory threshold (mm) | 13.31 ± 0.54 | 12.88 ± 0.61 | 13.82 ± 0.93 | 0.87 |
| Pain threshold (mm) | 32.10 ± 1.24 | 32.77 ± 1.63 | 31.32 ± 1.90 | 0.58 |
| Δ CS<sup>+</sup> valence | 68.08 ± 6.05 | 86.44 ± 7.04 | 45.94 ± 9.03 | <.001*** |
| Δ CS<sup>-</sup> valence | -23.47 ± 5.96 | -44.54 ± 6.62 | 1.94 ± 8.71 | <.001*** |
| CS<sup>+</sup> contingency accuracy (%) | 79.89 ± 2.49 | 88.65 ± 1.61 | 69.33 ± 4.54 | <.001*** |
| CS<sup>-</sup> contingency accuracy (%) | 78.89 ± 2.98 | 94.44 ± 1.30 | 60.15 ± 4.69 | <.001*** |

Descriptive statistics and results of independent samples t-tests comparing the low and high contingency
awareness group with respect to sociodemographic and psychological characteristics, sensory and pain thresholds, as well as behavioral markers of associative learning. All data are given as mean ± standard error of the mean. \*\*\* $p < .001$ , \*\* $p < .01$ , \* $p < .05$ . BMI body mass index, CS conditioned stimulus, HADS Hospital Anxiety and Depression Scale.

### 3.1 Modulation of CS-induced activation by contingency awareness

Results from the full factorial model to estimate the putative modulation of CS-induced activation patterns by contingency awareness revealed a significant main effect of CS, as previously reported in the respective subsamples (Icenhour et al., 2015a; Labrenz et al., 2016). Notably, a significant interaction between CS and contingency awareness group emerged in clusters of the dlPFC (*k_E_* = 110; *F* = 19.51; *p_FWE_* = .011) and parahippocampus (*k_E_* = 32; *F* = 14.54; *p_FWE_* = .029) (Figure 1A). Extracted contrast estimates supported very similar patterns for both regions, with a significant difference between CS^+^ and CS^-^ for dlPFC (*t*_40_ = 3.82, *p* < .001, *d* = .60) and parahippocampus (*t*_40_ = 3.81, *p* < .001, *d* = .59) exclusively in the high awareness group. In contrast, no significant differentiation was found in the low contingency group (both p > .351) Group effects were mainly driven by differences in CS^+^-induced activation of dlPFC (t_73_ = 2.47, *p* = .016, *d* = .57) and parahippocampus (*t*_73_ = 2.16, *p* = .034, *d* = .50), but not CS^-^-related differences (both *p* > .093).

**Figure 1.**
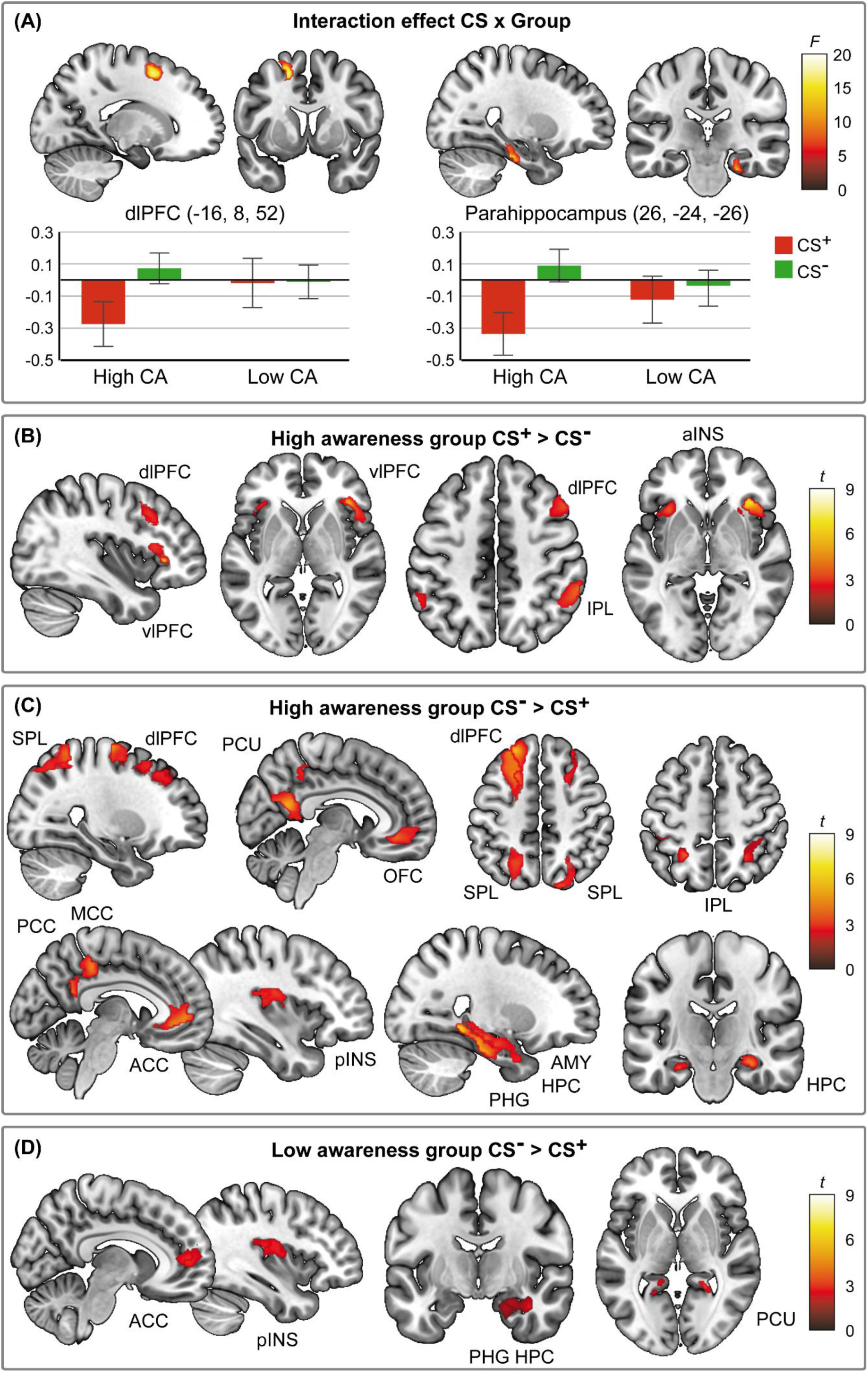
Differential neural activation patterns from the interaction between contingency awareness groups and CS. Results of the full factorial model depicting the interaction effect between contingency awareness (CA) groups and CS type with beta weights estimates indicating the direction of effects (A) and post-hoc paired t-tests for the high contingency awareness group with contrasts CS^+^ > CS^-^ (B) and CS^-^ > CS^+^ (C) as well as for the low contingency awareness group with the contrast CS^-^ > CS^+^ (D). Activations were superimposed on a structural T1-weighted MRI and masks for relevant ROI were applied for visualization purposes; color bars indicate respective *F*- and *t*-scores. For statistical details see Table 2. ACC anterior cingulate cortex, aINS anterior insula, AMY amygdala, dlPFC dorsolateral prefrontal cortex, HPC hippocampus, IPL inferior parietal lobule, MCC mid cingulate cortex, OFC orbitofrontal cortex, PCC posterior cingulate cortex, PCU precuneus, PHG parahippocampus, pINS posterior insula, SPL superior parietal lobule, vlPFC ventrolateral prefrontal cortex.

Post hoc tests were conducted to compare differential CS-induced activations within each group separately. The results obtained are summarized in Table 2 and illustrated in Figures 1B-D. Briefly, for the contrast CS^+^ > CS^-^, the high awareness group showed enhanced activation in the vlPFC, dlPFC, anterior insula, and IPL (Figure 1B).

**Table 2.** Differential CS-induced neural activation within contingency awareness groups.

| Brain region | H | Coordinates | | | $k_E$ | $t$ | $p_{FWE}$ |
| --- | --- | --- | --- | --- | --- | --- | --- |
|  |  | x | y | z |  |  |  |
| **High awareness CS<sup>+</sup> > CS<sup>-</sup>** |  |  |  |  |  |  |  |
| vlPFC (inferior frontal gyrus) | L | -36 | 22 | 8 | 48 | 4.18 | .013 |
|  | R | 34 | 30 | 2 | 361 | 5.97 | < .001 |
| dIPFC (middle frontal gyrus) | R | 52 | 20 | 44 | 234 | 4.67 | .003 |
| Anterior insula | L | -34 | 24 | 6 | 182 | 4.57 | .001 |
|  | R | 34 | 28 | 0 | 325 | 6.54 | < .001 |
| IPL | L | -54 | -50 | 40 | 87 | 3.69 | .035 |
|  | R | 56 | -48 | 50 | 212 | 5.08 | < .001 |
| **Low awareness CS<sup>+</sup> > CS<sup>-</sup>** |  |  |  |  | No significant clusters |  |  |
| **High awareness CS<sup>-</sup> > CS<sup>+</sup>** |  |  |  |  |  |  |  |
| dlPFC (middle frontal gyrus) | L | -26 | 36 | 46 | 801 | 5.53 | < .001 |
|  | R | 22 | 32 | 42 | 155 | 4.56 | .007 |
| dlPFC (superior frontal gyrus) | L | -22 | 38 | 50 | 1295 | 5.85 | < .001 |
|  | R | 14 | -16 | 72 | 397 | 5.90 | < .001 |
| OFC (orbital gyrus medial) | L | -8 | 38 | -10 | 360 | 4.76 | < .001 |
|  | R | 4 | 38 | -8 | 330 | 4.75 | < .001 |
| ACC | L | -10 | 48 | 2 | 474 | 5.52 | < .001 |
|  | R | 8 | 52 | 8 | 243 | 5.14 | < .001 |
| MCC | L | -2 | -36 | 46 | 248 | 4.40 | .003 |
|  | R | 2 | -32 | 52 | 168 | 4.30 | .005 |
| PCC | L | 0 | -54 | 22 | 109 | 5.24 | < .001 |
|  | R | 4 | -50 | 24 | 37 | 4.32 | .001 |
| Posterior insula | L | -36 | -8 | 4 | 29 | 4.05 | .005 |
|  | R | 38 | -10 | 20 | 154 | 4.66 | < .001 |
| IPL | L | -24 | -52 | 54 | 31 | 5.12 | < .001 |
|  | R | 26 | -52 | 54 | 27 | 4.42 | .002 |
| SPL | L | -26 | -52 | 60 | 860 | 5.33 | < .001 |
|  | R | 22 | -54 | 60 | 690 | 5.78 | < .001 |
| Precuneus | L | -6 | -40 | 74 | 381 | 5.84 | < .001 |
|  | R | 8 | -58 | 18 | 687 | 5.74 | < .001 |
| Amygdala | R | 28 | 0 | -24 | 49 | 3.85 | .003 |
| Hippocampus | L | -26 | -20 | -20 | 94 | 4.36 | .002 |
|  | R | 30 | -24 | -14 | 283 | 5.36 | < .001 |
| Parahippocampus | L | -26 | -42 | -8 | 322 | 5.56 | < .001 |
|  | R | 24 | -42 | -10 | 688 | 6.87 | < .001 |
| **Low awareness CS<sup>-</sup> > CS<sup>+</sup>** |  |  |  |  |  |  |  |
| ACC | L | -8 | 36 | 0 | 31 | 4.17 | .005 |
|  | R | 8 | 50 | 8 | 54 | 4.13 | .005 |
| Posterior insula | R | 34 | -8 | 16 | 143 | 4.15 | .003 |
| Precuneus | L | -24 | -58 | 4 | 9 | 3.92 | .025 |
|  | R | 24 | -48 | 0 | 21 | 4.91 | .001 |
| Hippocampus | R | 38 | -8 | -22 | 7 | 3.56 | .027 |
| Parahippocampus | R | 22 | -44 | -4 | 38 | 4.60 | .001 |

Results of region-of-interest analyses from paired t-tests comparing CS<sup>+</sup> > CS<sup>-</sup> and CS<sup>-</sup> > CS<sup>+</sup> within each contingency awareness group. Results of voxel-based analyses are provided with peak MNI coordinates and exact unilateral *p* values (all *p*<sub>FWE</sub> < .05). H hemisphere, *k*<sub>E</sub> cluster size, L left, R right.

At the same time, no significant results were found for the low awareness group. The reverse contrast (CS^-^ > CS^+^) revealed neural responses in a widespread network encompassing the vlPFC, dlPFC, OFC, cingulate and insular cortices, IPL, SPL, precuneus, amygdala, hippocampus, and parahippocampus in the highly aware group (Figure 1C). For the low awareness group, significant activation clusters were observed in the dlPFC, ACC, posterior insula, precuneus, hippocampus, and parahippocampus (Figure 1D).

### 3.2 Multiple regression analyses of CS-related activation and contingency awareness

Multiple regression analyses in the full sample were calculated to investigate CS-induced neural activation as a function of contingency awareness and CS valence change (Figure 2). While the effect of interest for the CS^+^ did not reveal significant activation clusters, for the CS^-^ enhanced neural responses were observed in the vlPFC, anterior insula, and IPL (Table 3). Post hoc test revealed this effect to be related to increased contingency awareness and an increase in positive CS^-^-valence (Table 4).

**Figure 2.**
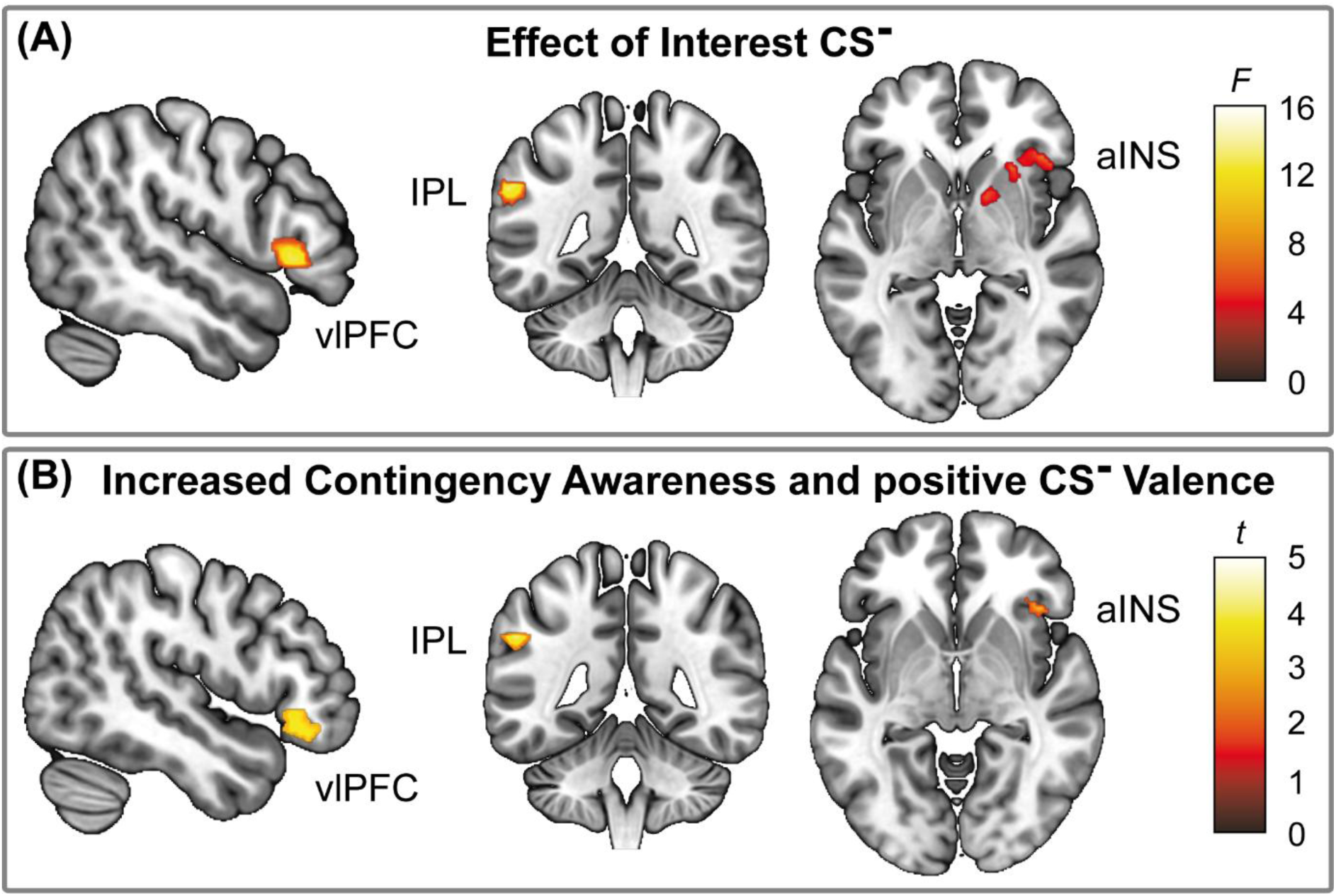
Results of linear regression analyses for the CS^-^. Results of linear regression analyses with covariates of contingency accuracy and changes in CS^-^ valence depicting for the full sample the Effect of Interest (A) and post hoc test for CS^-^ (B). Activations were superimposed on a structural T1-weighted MRI and masks for relevant ROI were applied for visualization purposes; color bars indicate respective F- and t-scores. For statistical details see Tables 3 and 4. aINS anterior insula, IPL inferior parietal lobule, vlPFC ventrolateral prefrontal cortex.

**Table 3.** Effects of Interest from multiple regression analyses.

| Brain region | H | Coordinates | | | $k_E$ | F | $p_{FWE}$ |
| --- | --- | --- | --- | --- | --- | --- | --- |
|  |  | x | y | z |  |  |  |
| CS<sup>+</sup> |  |  |  |  |  |  | No significant clusters |
| CS<sup>-</sup> |  |  |  |  |  |  |  |
| vIPFC (inferior frontal gyrus) | R | 52 | 22 | 2 | 173 | 14.14 | .006 |
| Anterior insula | R | 42 | 26 | -4 | 42 | 10.23 | .021 |
| IPL | L | -56 | -44 | 36 | 141 | 15.67 | .001 |

Region-of-interest results for Effects of Interest from multiple regression analyses including covariates of contingency accuracy and CS<sup>-</sup> valence change from baseline to post-acquisition. Results of voxel-based analyses are provided with peak MNI coordinates and exact unilateral *p* values (all *p*<sub>FWE</sub> < .05). H hemisphere, *k*<sub>E</sub> cluster size, L left, R right.

**Table 4.** Regression results for prediction of contingency awareness and positive CS^-^ valence.

| Brain region | H | Coordinates |  |  | *k*<sub>E</sub> | t | *p*<sub>FWE</sub> |
| --- | --- | --- | --- | --- | --- | --- | --- |
|  |  | x | y | z |  |  |  |
| vIPFC (inferior frontal gyrus) | L | -42 | 36 | -18 | 98 | 4.10 | .011 |
|  | R | 38 | 32 | -8 | 264 | 4.19 | .009 |
| Anterior insula | R | 42 | 26 | -4 | 8 | 3.49 | .046 |
| IPL | L | -54 | -44 | 34 | 40 | 4.72 | .001 |

Results for post-hoc test from multiple regression analyses reporting region-of-interest analyses of increasing contingency awareness and positive CS<sup>-</sup> valence. Results of voxel-based analyses are provided with peak MNI coordinates and exact unilateral *p* values (all *p*<sub>FWE</sub> < .05). H hemisphere, *k*<sub>E</sub> cluster size, L left, R right.

### 3.3 Pain-related activation

Exploratory analyses were additionally conducted to evaluate whether stronger CS-related differentiation in highly aware individuals could be related to group differences in US-related neural activation. Comparisons of high vs. low contingency awareness groups did not reveal any differences in the central processing of visceral pain.

## 4 Discussion

The significance of pain-related fear in the pathophysiology and treatment of chronic pain is increasingly acknowledged, prompting experimental research to uncover mechanisms underlying learning and memory processes in the context of pain (Elsenbruch et al., 2020; Labrenz et al., 2023, 2022b; Meulders, 2020; Vlaeyen, 2015). Human fear conditioning studies using highly aversive unconditioned stimuli indicate distinct emotional learning processes, characterized not only by negative emotions in response to cues signaling danger to the organism but also by positively valued responses to safety signals as conditioned inhibitors of fear (Christianson et al., 2012; Labrenz et al., 2022b; Laing et al., 2021). Behaviorally, these processes manifest as changes in the perceived valence of previously neutral predictive cues, but are likely also mediated by distinct neural networks (Fullana et al., 2016), which seem to be altered in chronic pain patients (Claassen et al., 2017; Icenhour et al., 2015b; Lanters et al., 2024; Meier et al., 2018). An important, yet unresolved question remains the potential role of conscious awareness of cue-outcome relationships in influencing the formation of emotional memories, particularly in the context of visceral pain along the gut-brain axis.

We therefore investigated the neural correlates of contingency awareness in shaping fear and safety learning in an experimental model of visceral pain. Our findings provide first evidence in the field of interoceptive visceral pain that a pronounced differentiation in the neural processing of pain- and safety-predictive cues during associative learning requires cognitive awareness. They further highlight that contingency awareness is particularly relevant in encoding and associating safety-rather than threat-related environmental information. These findings on a neural level complement and extend our earlier behavioral observations of more pronounced negative valence in response to the CS^+^ and stronger positive emotions in response to the CS^-^ following pain-related fear learning in highly aware compared to unaware individuals (Labrenz et al., 2015a).

Specifically, a learned differentiation between threat and safety cues manifested as a function of accurate cognitive awareness of cue-pain relations, with highly aware individuals exhibiting substantially more pronounced differential responses of dlPFC and parahippocampus as core regions of the emotional arousal and executive control networks. Via projections to other prefrontal, as well as parietal, and limbic regions (Ma et al., 2022; van Strien et al., 2009), this key hub is tightly linked to various cognitive and emotional functions, particularly affective working memory, emotional memory formation and emotion regulation (Aminoff et al., 2013; Dolcos et al., 2012; Dolcos & Denkova, 2014; Kensinger & Corkin, 2004; Li et al., 2016; Mikels & Reuter-Lorenz, 2019), highlighting the central role in the formation of cognitive awareness by enabling an accurate detection, processing, and emotional differentiation of associations relevant to threat and safety.

The link between these (para)limbic and prefrontal regions has previously been discussed, indicating that emotions can selectively strengthen or weaken memory traces via direct limbic input or by indirect top-down attentional control (Salsano et al., 2024; Vuilleumier, 2005). Our findings support an interplay between both mechanisms in the processing of signals depending on their value as threat- or safety-relevant.

To substantiate these findings, we evaluated threat- and safety-related neural activation separately in their predictive value for cognitive and emotional correlates of visceral pain-related learning. Neural activation during visceral pain anticipation was not predictive of accurate contingency awareness or changes in negative emotional valence on a behavioral level, in line with our previous observations (Labrenz et al., 2015a). Conversely, the processing of conditioned safety cues recruited substantial resources encompassing the fronto-parietal network in concert with cingulate, insular, and limbic regions, and predicted accurate contingency and positive valence. These activation patterns fit neatly into existing literature on safety-related processing (Christianson et al., 2012; Fullana et al., 2016) and our previous insights from an experimental study, in which we manipulated the possibility to establish contingency awareness by conditioning a group with predictable and one with unpredictable visceral pain (Labrenz et al., 2016). We observed enhanced differential neural activation in regions of the emotional arousal and executive control networks in the group able to predict pain or safety from it, which was mainly attributable to differences in the processing of contingently learned safety cue properties. Together, these results suggest that putatively different processes underlie the acquisition and inhibition of fear.

On the one hand, fear acquisition may occur through an evolutionarily hard-wired and therefore cognitively less demanding mechanism that enables individuals to immediately prepare for emerging threats (Seligman, 1971; Öhman, 2005). In contrast, fear inhibition processes such as differentiating between reinforced and non-reinforced stimuli, extinction, or conditioned inhibition, which involve learning about safety cue properties, may depend on cognitive models and therefore require contingency awareness (Jovanovic et al., 2006; Lovibond, 2004), with crucial clinical implications for exposure-based treatments.

Together, our findings strongly advocate for the distinct consideration of conditioned responses to pain-related threat and safety cues. Relying solely on differential measures, a common practice in fear conditioning studies, may obscure their unique neural correlates (Fullana et al., 2016) that appear to be particularly relevant to visceral pain-related emotional learning and memory processes (Benson et al., 2014; Gramsch et al., 2014; Icenhour et al., 2015a, b; Labrenz et al., 2022a, 2015b). Furthermore, the ongoing debate about the potential impact of contingency awareness has often overlooked safety learning (Klucken et al., 2009; Lovibond et al., 2011; Schultz & Helmstetter, 2010; Tabbert et al., 2011). Consequently, in the controversy of whether cognitive awareness is non-essential or rather a prerequisite for successful emotional memory encoding, both views may be valid.

Finally, our findings support previous assertions that safety-related learning processes warrant greater attention in the context of pain (Vlaeyen, 2015) and the broader field of fear conditioning (Christianson et al., 2012; Fullana et al., 2016). Considering evidence that contingency learning is altered in chronic pain (Icenhour et al., 2015b; Jenewein et al., 2013; Meulders et al., 2014), our results underscore the need for further investigation to elucidate the potential interplay between attention and contingency awareness in impaired extinction and its relevance to exposure-based interventions in patients with chronic pain.

Recognizing how individuals perceive and respond to threat and safety cues can improve therapeutic approaches and foster personalized treatments to strengthen accurate safety processing and reduce maladaptive fear responses. For instance, cognitive-behavioral therapy and mindfulness-based interventions can help patients reframe their awareness and reduce the psychological impact of visceral pain (Lackner et al., 2007) Educating patients about the gut-brain axis and the role of contingency awareness and providing them with knowledge about the mechanisms underlying their pain can reduce anxiety and maladaptive avoidance (Labrenz et al., 2022b), enable them to manage their symptoms more effectively, and improve treatment adherence (Keefer et al., 2022).

Our experimental findings, derived from a model of acute visceral pain in a sample of young and healthy individuals, undoubtedly carry limited generalizability to affected patient populations, adding only a small piece to the puzzle. Ultimately, however, recognizing the role of contingency awareness and underlying mechanisms offers a promising path toward refining exposure-based therapies to enhance treatment efficacy in disorders of gut-brain interaction.

## Research Data Availability Statement

The data that support the findings of this study are not openly available but are available from the corresponding author upon reasonable request. Data are located in controlled access data storage at Ruhr University Bochum.

## Acknowledgements

We would like to thank Dr. Nikolai Axmacher and Dr. Erhan Genç for excellent support in data analyses.

